# Let’s face it: The lateralization of the face perception network as measured with fMRI is not clearly right dominant

**DOI:** 10.1101/2022.02.06.479156

**Authors:** Ina Thome, José C. García Alanis, Jannika Volk, Christoph Vogelbacher, Olaf Steinsträter, Andreas Jansen

## Abstract

The neural face perception network is distributed across both hemispheres. However, the dominant role in humans is virtually unanimously attributed to the right hemisphere. Interestingly, there are, to our knowledge, no imaging studies that systematically describe the distribution of hemispheric lateralization in the core system of face perception across subjects in large cohorts so far. To address this, we determined the hemispheric lateralization of all core system regions (i.e., occipital face area (OFA), fusiform face area (FFA), posterior superior temporal sulcus (pSTS)) in 108 healthy subjects using functional magnetic resonance imaging (fMRI). We were particularly interested in the variability of hemispheric lateralization across subjects and explored how many subjects can be classified as right-dominant based on the fMRI activation pattern. We further assessed lateralization differences between different regions of the core system and analyzed the influence of handedness and sex on the lateralization with a generalized mixed effects regression model. As expected, brain activity was on average stronger in right-hemispheric brain regions than in their left-hemispheric homologues. This asymmetry was, however, only weakly pronounced in comparison to other lateralized brain functions (such as language and spatial attention) and strongly varied between individuals. Only half of the subjects in the present study could be classified as right-hemispheric dominant. Additionally, we did not detect significant lateralization differences between core system regions. Our data did also not support a general leftward shift of hemispheric lateralization in left-handers. Only the interaction of handedness and sex in the FFA revealed that specifically left-handed men were significantly more left-lateralized compared to right-handed males. In essence, our fMRI data did not support a clear right-hemispheric dominance of the face perception network. Our findings thus ultimately question the dogma that the face perception network – as measured with fMRI – can be characterized as “typically right lateralized”.

## 1. INTRODUCTION

### 1.1. The neural network underlying face perception

Face perception is mediated by a distributed neural network. This network is, as proposed more than 20 years ago by Haxby and colleagues, often divided into a “core system” and an “extended system” (Haxby et al., 2000; Haxby and Gobbini, 2011). The core system is associated with the analysis of the visual appearance of faces. It consists of three bilateral brain regions in the occipito-temporal cortex: the occipital face area (OFA) in the inferior occipital gyrus, the fusiform face area (FFA) in the lateral fusiform gyrus and the posterior superior temporal sulcus (pSTS). Each of these regions has a distinct role in the process of face perception. The OFA is typically associated with the analysis of invariant facial features like eyes or mouth and helps to decide if an object is a face or not (Gauthier et al., 2000; Haxby et al., 1999; Pitcher et al., 2011b). The FFA predominantly processes identity (Kanwisher et al., 1997), while the pSTS engages in the extraction of changeable features such as expression, eye-gaze and lip movement (Engell and Haxby, 2007; Nummenmaa et al., 2010; Puce et al., 1998). The extended system is distributed over limbic, parietal and prefrontal regions. It is associated with the retrieval of person-knowledge and other nonvisual information. For example, the auditory cortex guides speech perception, the anterior temporal lobe is said to contribute semantic and biographic information and the insula and amygdala come into play, when emotional aspects are involved (Duchaine and Yovel, 2015; Haxby et al., 2000; Haxby and Gobbini, 2011). More recent studies reported additional face sensitive areas in the anterior temporal lobe (ATL) (Rajimehr et al., 2009; Tsao et al., 2008), the anterior superior temporal sulcus (aSTS) (Pitcher et al., 2011a) and the ventral lateral prefrontal cortex (often referred to as inferior frontal gyrus (IFG) or inferior frontal junction (IFJ)) (Chan and Downing, 2011; Pitcher et al., 2011a). Furthermore, Weiner and Grill-Spector (2012) questioned the idea of one single FFA and proposed a subdivision into a medial FFA (mFFA, located in medial fusiform gyrus) and a posterior FFA (pFFA, located in posterior fusiform gyrus) instead. All these discoveries inspired Duchaine and Yovel (2015) to propose a revised neural network for face perception. It includes all the aforementioned face sensitive brain areas and assigns them to a ventral (OFA, FFA and ATL) and a dorsal (pSTS, aSTS and IFG) pathway.

### 1.2. Hemispheric lateralization of the face perception network

The neural face perception network is distributed across both hemispheres. However, the dominant role in humans is virtually unanimously attributed to the right hemisphere. This finding first originated from lesion studies. Here, it has been observed that most patients suffering from acquired prosopagnosia, i.e., the inability to recognize the identity of faces following brain damage, had lesions in the right posterior hemisphere (for an overview, cf. Bukowski et al., 2013). In contrast, prosopagnosia following unilateral lesions to the left hemisphere has been reported only in few cases (Barton, 2008; Eimer and McCarthy, 1999; Mattson et al., 2000; Tzavaras et al., 1973). The right-hemispheric dominance of the face-processing network was subsequently confirmed in various other studies. It is now based on ample evidence accumulated over decades of research with lesion patients, brain stimulation techniques or behavioral experiments (for an overview, see Duchaine and Yovel, 2015; Rossion and Lochy, 2021).

Also the functional magnetic resonance imaging (fMRI) literature seems to confirm, at least at first glance, the right-hemispheric dominance of the face perception network. Even though brain activation in both hemispheres is reported for face processing tasks, the right hemisphere is usually described as “dominant”. It shows overall stronger responses to face stimuli, both in terms of the spatial extent of the activation and the strength of activity (Badzakova-Trajkov et al., 2010; Frässle et al., 2016c; Ishai et al., 2005; Rhodes et al., 2004; Rossion et al., 2012). Interestingly, a more thorough analysis of the literature provides a more ambiguous picture. More specifically, while fMRI studies that describe hemispheric lateralization across the averaged activation in the entire core system or in even larger regions (e.g., the entire temporal lobe) often report a clear right-hemispheric dominance (e.g., Badzakova-Trajkov et al., 2010), other studies that calculate lateralization for individual regions of the core system often report a high interindividual variability (e.g., Canário et al., 2020; Davies-Thompson et al., 2016; De Winter et al., 2015; Johnstone et al., 2020). This high variability results in up to 45% of subjects being not right-hemispheric dominant for face perception.

### 1.3. Interindividual variabiliy of hemispheric lateralization

High interindividual variability of the hemispheric lateralization of the face peerception network is also in accordance with our own anecdotal experience. Our research group has conducted numerous fMRI studies on face processing over the last years, often in the context of hemispheric lateralization (e.g., Frässle et al., 2016a, 2016b, 2016c; Hildesheim et al., 2020; Sahraei et al., 2021; Thome et al., 2021; Zimmermann et al., 2019). In these studies, it was often necessary to assess the fMRI activation patterns not only at the group level, but also at the individual level, e.g., in order to determine the spatial location of core system regions for further analyses. Here we noticed, independent of the specific face processing task, that although the face network was consistently (albeit not strongly) right-lateralized at the group level, there was a strong variability at the individual level. Even among right-handers, many subjects showed a bilateral or left-hemispheric lateralization. So far, however, we never assessed the distribution of hemispheric lateralization of the face perception network systematically. Interestingly, there are, to our knowledge, also no other imaging studies yet that investigated the interindividual variation of hemispheric lateralization of face perception across subjects in large cohorts. This is in clear contrast to the investigation of for instance the language or the spatial attention network (e.g., Flöel et al., 2005; Jansen et al., 2007; Knecht et al., 2000a, 2000b; Springer et al., 1999). It is thus unknown how many individuals can be characterized as right-dominant, left-dominant or bilateral for face perception based on the fMRI activation pattern. <u>The first aim of the present study</u> was therefore to thoroughly decribe hemispheric lateralization of all regions of the core system of face perception in a large cohort of subjects (including left- and right-handers). We aimed to assess the variability of hemispheric dominance within the population and to determine to which degree the network is lateralized to the right hemisphere based on the fMRI activation pattern.

### 1.4. Effects of region, handedness, and sex on hemispheric lateralization

The hemispheric lateralization of cognitive functions is in general highly flexible and can be modulated by various factors like handedness, sex, age, genetic factors, hormonal influences, or disease (Toga and Thompson, 2003). For example, in language research a relationship between handedness and hemispheric dominance is well established. While 96% of strong right-handed subjects show a left hemispheric language dominance, this value is reduced to 85% in ambidextrous individuals and 73% in strong left-handers (Knecht et al., 2000b). In a similar vein, a number of neuroimaging studies reported a relationship between handedness and hemispheric lateralization also for face perception. Willems et al. (2010) showed that the typical right-ward lateralization of the FFA was absent in left-handers who, on average, showed a more bilateral activation pattern. Bukowski et al. (2013) replicated these findings and additionally showed that this reduced right-hemispheric lateralization was specific to the FFA, while OFA and STS were right-lateralized in both right- and left-handers, without a difference in the degree of lateralization. A possible explanation for this spatial specifity is often attributed to the left dominance of the visusal word form area (VWFA), a region associated with the identification of words and letters from lower-level shape images, prior to association with phonology or semantics (Dehaene and Cohen, 2011; Price and Devlin, 2003; for an overview also see Hildesheim et al., 2020; Rossion and Lochy, 2021). In another recent study, Frässle et al. (2016a) combined fMRI and Dynamic Causal Modeling (DCM) to elucidate the neural mechanisms underlying the different hemispheric lateralization of face perception in right- and left-handers. They reported an enhanced recruitment of the left FFA in left-handers, as shown by stronger face-specific modulatory influences on both intra- and interhemispheric connections.

<u>The second aim of the present study</u> was to explore the influence of different factors on hemispheric lateralization of face perception. More specifically, we aimed to investigate effects of region (OFA, FFA, STS), handedness, and sex on the degree of lateralization. We expected, as outlined above, a reduced right-hemispheric lateralization of the FFA in left-handers compared to right-handers (Bukowski et al., 2013; Frässle et al., 2016a; Willems et al., 2010). We further explored whether the OFA, often considered to be a hierarchically lower region of the face perception network, is characterized by decreased lateralization compared to the FFA and STS (Rossion et al., 2012).

## 2. MATERIALS AND METHODS

### 2.1. Subjects

Subjects were recruited in an ongoing fMRI study investigating the neural mechanisms underlying hemispheric lateralization. At the time of data analysis, 119 subjects had been included. Nine subjects had to be excluded due to bad quality of MRI data. Two subjects were excluded because they could not be clearly classified as either right- or left-hander. One-hundred-eight subjects (67 females, 41 males; mean age 24.5 years ± 3.6 years) were therefore included in the final data analysis (see demographics in Table 1). Eighty-five participants were right-handed and twenty-three left-handed, as assessed by the twelve-item short version of the Edinburgh Handedness Inventory (Oldfield 1971, cut-off +/-30). The proportion of left-handed subjects (~21%) was higher than would have been expected if the sample had been randomly selected (~10%; Coren and Porac 1977; McManus 2019). However, we deliberately chose to increase the proportion of left-handers in order to explore the effect of handedness on hemispheric lateralization. All subjects had normal or corrected-to-normal vision and no history of neurological or psychiatric disorders. Written consent to participate in the study was given by all participants.

**Table 1.** Cohort demographics (mean age and standard deviation [SD] in brackets)

|  | All | Female | Male |
| --- | --- | --- | --- |
| All subjects | 108 (24.5, SD = 3.6) | 67 (23.4, SD = 2.9) | 41 (26.2, SD = 4.0) |
| Right-handers | 85 (24.1, SD = 3.2) | 51 (23.1, SD = 2.7) | 34 (25.6, SD = 3.4) |
| Left-handers | 23 (25.9, SD = 4.6) | 16 (24.5, SD = 3.2) | 7 (29.1, SD = 5.7) |

The experiment was implemented in accordance with the Declaration of Helsinki and was approved by the local ethics committee of the Medical Faculty of the University of Marburg, where all imaging took place (file reference 160/13 version 2).

### 2.2. Experimental paradigm

Subjects viewed either faces, houses or scrambled images in a blocked design. All stimuli were presented using the Presentation software 18.1 (Neurobehavioral Systems Inc., Berkeley, California, United States, 2000). The scrambled images were generated by applying a Fourier transformation to the original images of both faces and houses. Face stimuli were taken from the Centre for Vital Longevity Face Database (Ebner, 2008), the house stimuli were kindly provided by Joshua Goh (Goh et al., 2010). All images were static, frontal, black and white photographs. The stimuli were presented centrally. During the whole paradigm, a fixation cross was shown in the middle of the screen. Participants were instructed to fixate this cross and perceive the stimuli around it. Nine blocks of each stimulus category were presented in pseudo-randomized order, each containing 20 stimuli. Stimuli were presented for 300 ms and were followed by a fixation cross for 425 ms. Each block lasted for 14.5 seconds. Stimulus blocks were separated by baseline blocks of 14.5 seconds, where only the fixation cross was shown. In the middle of the experiment, there was a short pause of 20 seconds. This resulted in a total length of approximately 13 minutes for the whole face processing task. To ensure that participants were paying attention during this passive viewing task, they were instructed to always press a button when the same image appeared twice in a row (1-back task).

### 2.3. MRI data acquisition

Subjects were scanned on a 3 Tesla MR scanner (TIM Trio, Siemens, Erlangen, Germany) with a 12-channel head matrix receive coil at the Department of Psychiatry, University of Marburg. Functional MRI images were acquired with a T2*-weighted gradient-echo echo planar imaging sequence sensitive to the blood-oxygenation-level-dependent (BOLD) contrast (TR = 1450 ms, TE = 25 ms, voxel size = 3×3×4 mm^3^, 30 slices, 4 mm thickness, flip angle = 90°, matrix size = 64 × 64 voxels, FoV = 192 × 192 mm^2^). Slices were measured in interleaved descending order parallel to the intercommissural plane (anterior to posterior commissure).

For each subject, a high-resolution T1-weighted anatomical image was collected using a magnetization-prepared rapid gradient-echo (3D MP-RAGE) sequence in sagittal orientation (TR = 1900 ms, TE = 2.54 ms, voxel size = 1 × 1 × 1 mm^3^, 176 slices, 1 mm thickness, flip angle 9°, matrix size = 384 × 384, FoV = 384 × 384 mm).

### 2.4. MRI data analysis

#### Preprocessing

Pre-processing was conducted using SPM12 (Statistical Parametric Mapping, version v6015, Wellcome Trust Centre for Neuroimaging, London, UK; http://www.fil.ion.ucl.ac.uk) and MATLAB R2009b (MathWorks, Natick, MA, USA) with an in-house pipeline that consisted of the following steps: realignment, coregistration, segmentation, normalization, and smoothing.

After discarding the first four functional scans which are prone to magnetization instability artefacts, all remaining functional images were corrected for head motion (realignment). The six realignment parameters were saved for further analyses. The individual images were realigned to the mean image and afterwards co-registered with the high-resolution anatomical T1-weighted image. Normalization to MNI (Montreal Neurological Institute) space was conducted using the segmentation-normalization approach (Ashburner and Friston, 2005). During this normalization step, the functional images were resampled to a voxel size of 2 × 2 × 2 mm^3^. After that, the normalized functional images were spatially smoothed with a 6 mm full width at half maximum (FWHM)-Gaussian kernel.

#### Statistical analysis

A first-level analysis for every subject’s functional data was conducted using a General Linear Model (GLM; (Friston et al., 1995). We modelled each condition (“faces”, “houses”, “scrambled images”) as a regressor, convolved with the canonical hemodynamic response function implemented in SPM. Furthermore, to control for movement-related artifacts, the six realignment parameters were introduced as nuisance regressors. Low-frequency noise in the data was accounted for by a high-pass filter (cut-off frequency: 1/128 Hz).

Individual brain activation in the core network was assessed by means of a conjunction analysis. With the conjunction, one is able to control both high- and low-level visual characteristics of faces (Rossion et al., 2012). Here, we first calculated contrast images and t-statistic images for the contrasts “faces > houses” and “faces > scrambled”. The conjunction t-map was then calculated by choosing for each voxel the smallest t-value from the “faces > houses” contrast and the “faces > scrambled” contrast (minimum statistic approach as suggested by Nichols et al. 2005). The resulting conjunction t-map (i.e., conjunction null hypothesis) provides a more conservative indicator of face-sensitivity compared to the conjunction analysis implemented in SPM (i.e., global null hypothesis; Friston et al., 2005).

At the group level, the individual contrast images were entered in a random-effects analysis. We specified a one-way ANOVA with two levels. For each level, we chose the contrast images either form the “faces > houses” contrast or the “faces > scrambled” contrast. We defined one contrast for each level (i.e., using the weights 1 0 and 0 1, respectively). Face sensitive activation was analyzed with a conjunction of those contrasts.

### 2.5. Assessment of hemispheric lateralization

Hemispheric lateralization for a specific cognitive task can be described by a lateralization index (LI). The LI, sometimes also referred to as asymmetry index (Anderson et al., 2006), quantifies whether the brain activation is predominantly left-hemispheric, right-hemispheric or bilateral. The LI is typically calculated with the following formula (among others, Binder et al. 1996; Jansen et al. 2006):

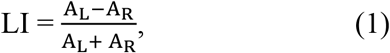

where A_L_ and A_R_ quantify the strength of fMRI-measured activity (A) within regions of interest (ROIs) of the left (L) and right (R) hemisphere, respectively. It results in an LI value ranging from −1 to +1. Negative values indicate a right-hemispheric dominance and positive values indicate a left-hemispheric dominance. The cut-off for bilateral activation can be set arbitrarily. However, as it is typically set to ± 0.2 in many studies (Bradshaw et al., 2017), we decided to also use this cut-off in the present study.

#### Choice of activity measure

In lateralization research several approaches have been established to quantify the strength of brain activity (i.e., A_L_ and A_R_). The most widely used measures of brain activity are either based on the magnitude of the fMRI signal change (weighted β-values or t-values) or the extent of the activated brain region (i.e., number of activated voxel) (see Jansen et al. (2006) for a detailed overview). In the present study, we used the magnitude of signal change defined by the t-values as activity measure. All LIs were calculated using the bootstrap procedure (Wilke and Schmithorst, 2006) implemented in the LI toolbox (version 1.3, Wilke and Lidzba 2007) for SPM12 (MATLAB version 2017a). This calculation encompassed the following steps: First, the individual conjunction t-maps were thresholded and masked with custom ROI-masks for the three core system regions (for creation of ROI masks, see below). Second, from all surviving voxels 100 bootstrapped samples were drawn from each hemisphere (resampling ration *k* = 0.25, with replacement) and all possible LI combinations (10,000) were calculated and plotted in a histogram. Third, from the central 50% of LI values a “trimmed mean” LI value was calculated. This procedure was repeated for all 20 regularly spaced thresholding steps. Finally, a weighted-overall mean was calculated by applying the t-threshold as a weighting factor. Hence, statistically more conservative thresholds lead to progressively stronger weightings. A more detailed description of the bootstrapping approach and the toolbox is given in Wilke and Schmithorst (2006) and Wilke and Lidzba (2007) respectively.

#### Definition of regions of interest

Ideally, ROI masks should encompass the relevant brain activation (sensitivity) without including other activated clusters (specificity). ROIs can either be determined anatomically (i.e., based on anatomical landmarks) or functionally (i.e., based on the activation pattern). As the core system of face perception is comprised of at least three brain areas in each hemisphere that are in close anatomical proximity, we decided to use functionally determined ROIs.

The exact localization of face perception areas in the core system varies highly between individual subjects. Therefore, we did not use one ROI mask for all subjects, but built subject-specific masks using the following procedure: First, we created for each ROI symmetrical box-shaped masks with the WFU PickAtlas (Maldjian et al., 2004, 2003) (v 3.0.5). Center coordinates and spatial extent was based on typical locations for OFA, FFA and STS reported in previous fMRI studies using face localizers (FFA, OFA, STS: Fox et al., 2009; FFA: Berman et al. 2010; right OFA, right FFA, right STS: Pitcher et al., 2011b; OFA, right STS: Rossion et al., 2012) and a search on the automated meta-analysis platform Neurosynth.org (STS, neurosynth.org/analyses/terms/psts/). The coordinates are summarized on this studies Open Science Framework repository (OSF, https://osf.io/s8gwd/). These fairly large masks were used as anatomical restriction. The center of the subject-specific ROIs had to be within these masks (see below). Second, we assessed the brain activation pattern at the group level. For each ROI, peak voxels were identified for the group-level conjunction contrast at p < 0.05, family-wise error (FWE) corrected (Table 2). Third, for each subject, all local maxima of the single subject conjunction t-map were determined that (i) were within the borders of the anatomical mask (created in the first step), (ii) had a t-value of at least 3.1 (corresponding to p < 0.001 uncorrected), and (iii) had t-values at least as high as the t-value at the group maximum coordinate. Fourth, the nearest local maximum to the group maximum was identified. If no local maximum met these criteria, we used the coordinate at the group maximum for this subject. Fifth, custom sphere-shaped masks with a radius of 10 mm centered around these individually determined coordinates were created. All these steps were performed with custom MATLAB codes. The individual center coordinates for each ROI are depicted in Fig. 1 (purple: OFA, green: FFA, yellow: STS).

**Table 2.**
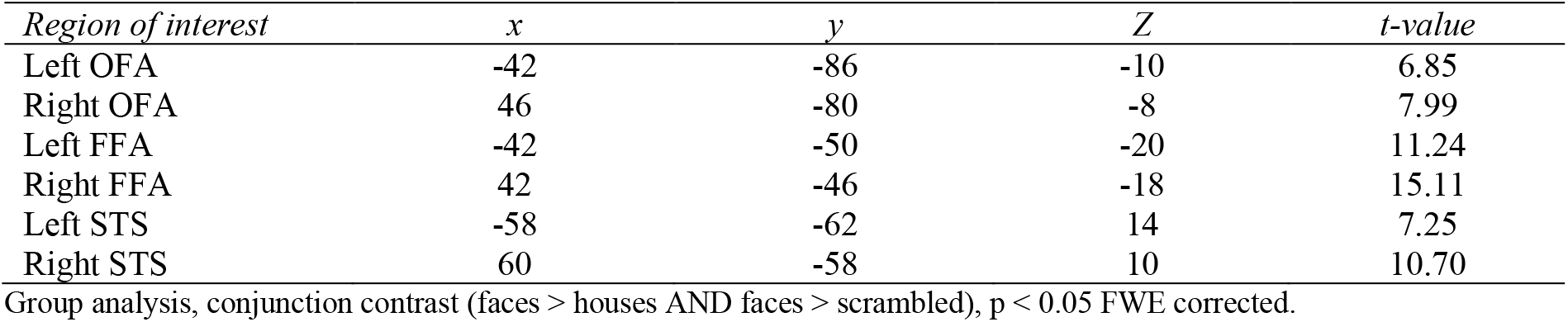
Core system MNI (x, y, z) coordinates.

| *Region of interest* | *x* | *y* | *Z* | *t-value* |
| --- | --- | --- | --- | --- |
| Left OFA | -42 | -86 | -10 | 6.85 |
| Right OFA | 46 | -80 | -8 | 7.99 |
| Left FFA | -42 | -50 | -20 | 11.24 |
| Right FFA | 42 | -46 | -18 | 15.11 |
| Left STS | -58 | -62 | 14 | 7.25 |
| Right STS | 60 | -58 | 10 | 10.70 |

Group analysis, conjunction contrast (faces > houses AND faces > scrambled), $p < 0.05$ FWE corrected.

**Fig. 1.**
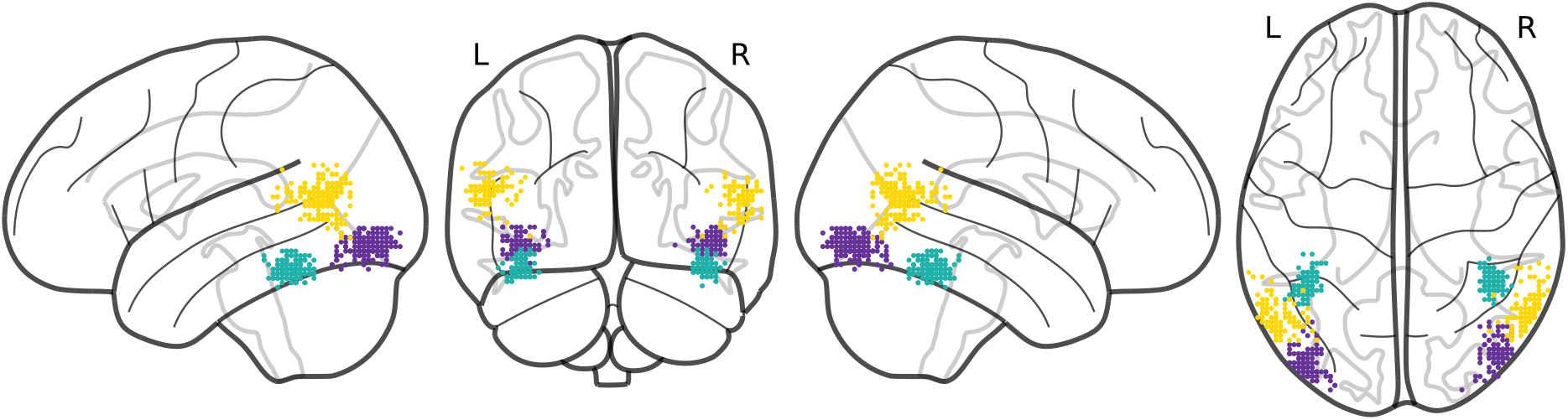
Individual maxima for the three brain regions of the core system of face perception: OFA (purple), FFA (green), STS (yellow). Coordinates were visualized with nilearn (version 0.7.0) (*Abraham et al., 2014*).

### 2.6. Effects of brain region, handedness, and sex on hemispheric lateralization

To assess the dependence of the LI on brain region (OFA, FFA, STS), handedness (right-handed, left-handed) and sex (male, female), we calculated the main effect for each factor as well as their interactions. All statistical analyses were carried out in the R programming environment (R Core Team, 2021, version 4.1.2). Data were analyzed using a generalized linear mixed effects regression approach assuming a gamma distribution and applying a log link function. These analyses were carried out using the glmer() function from the R-package lme4 (Bates et al., 2015). LI values were transformed with +1 in order to meet the prerequisites of a gamma distribution (i.e., positive-only). Like ordinary least squares (OLS) models, mixed effects regression examines the relationship between a set of predictors (e.g., brain region, handedness) and a response variable (e.g., LI-value). However, the repeated measures design (multiple measurements extracted from one subject) of the study might lead to strong interdependencies in the data, thus violating one of the key assumptions (the conditional mean should be zero) of OLS models (Ernst and Albers, 2017). To address this issue, we used a mixed models approach to account for individual variation of the response variable’s variance (e.g., more similar LI-values within subjects than between subjects), which, if led unaddressed, can lead to increased error variance in the ordinary regression models, diminishing their validity and statistical power. Furthermore, a mixed effects regression framework, allowed us to handle unbalanced data structures (i.e., due to missing data) more efficiently by nesting observations within subjects. All models were estimated via maximum likelihood. Main effects and interactions were assessed via Type III Wald Chi^2^-tests as implemented in R-package car (Fox and Weisberg, 2019). All categorical variables were effect (i.e., deviation) coded and all continuous variables mean centered around zero prior to analyses. Pairwise contrasts were computed on the basis of the estimated marginal means using the R-package emmeans (Lenth, 2021) and all p-values adjusted according to the false discovery rate (FDR; Benjamini and Hochberg 1995). Model descriptives, diagnostics, and estimates of effect sizes (standardized beta coefficients) are provided on OSF (https://osf.io/s8gwd/) along with the R-scripts and data to reproduce the analyses.

## 3. RESULTS

The results section is divided in three parts. First, we present the brain activation pattern for the face perception task (3.1.). Second, we describe the variability of hemispheric lateralization across subjects (3.2.). Last, we analyze the effects of region (OFA, FFA, STS), handedness and sex on hemispheric lateralization (3.3.).

### 3.1. Brain activation pattern associated with face perception

The face perception task was associated with brain activity in a distributed network encompassing the bilateral occipito-temporal cortex (including the core system’s brain regions OFA, FFA, and STS), frontal and parietal areas. For illustrative purposes, we present both the group activation pattern for right-handed subjects and an individual activation pattern for a selected right-handed subject in Fig. 2. On OSF (https://osf.io/s8gwd/) we additionally present the group activation pattern for all subjects and the group activation pattern for left-handed subjects.

**Fig. 2.**
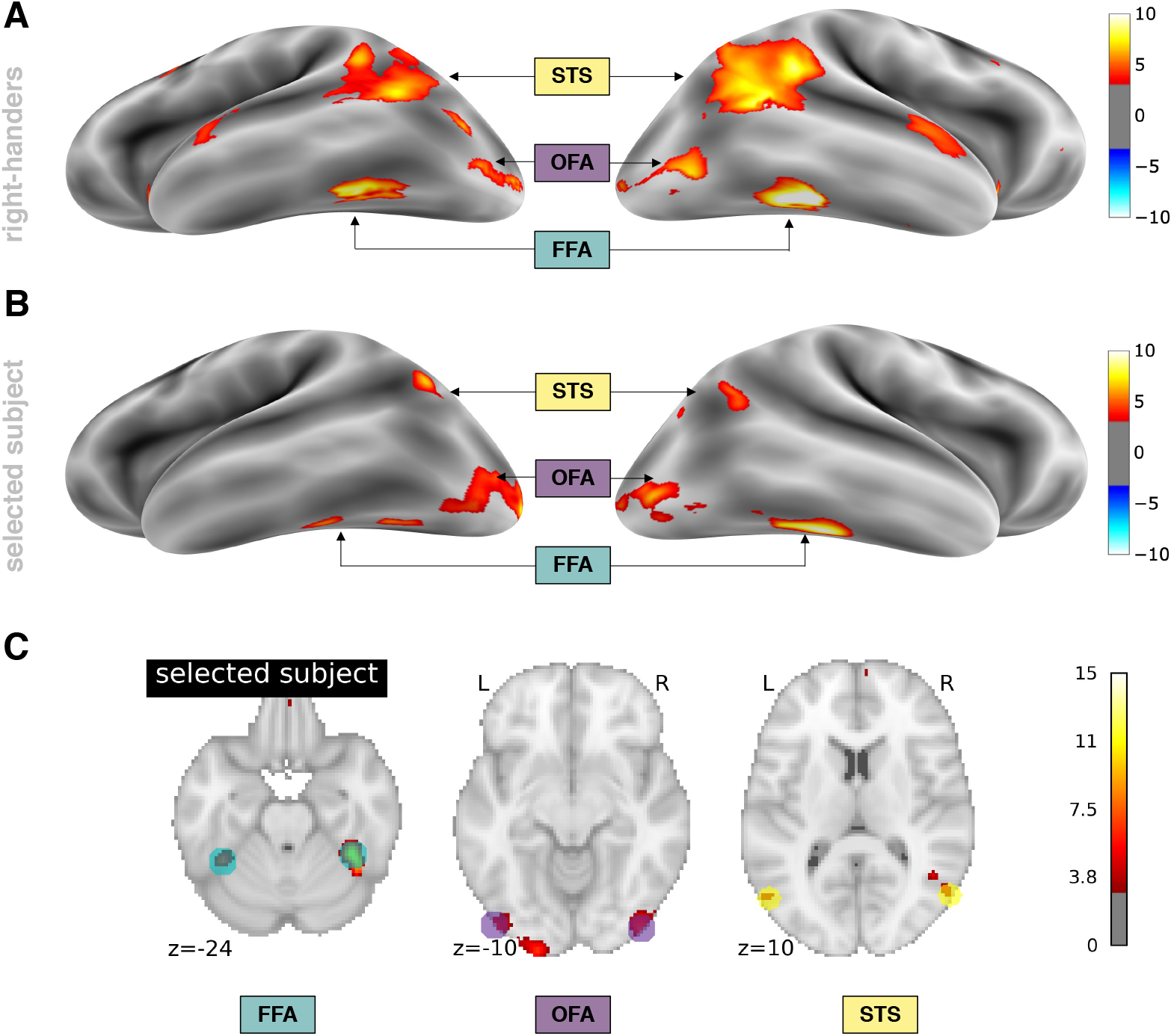
Brain activation during face perception. The activation pattern is assessed with the conjunction contrast “faces > houses” AND “faces > scrambled”. For illustrative purposes, we applied a statistical threshold of p = 0.001 (uncorrected). Activity in the core system of face perception (i.e., OFA, FFA, and STS) is clearly detectable. (**A**) The activation pattern for the group of right-handed subjects (n = 85) and (**B**) a selected subject’s brain activation (right-handed, female) is displayed on the inflated FreeSurfer fsaverage template (Fischl et al. 1999a; Fischl et al. 1999b). (**C**) Brain activity of the selected subject shown in B is additionally displayed on sections of the MNI ICBM152 T1 template (Fonov et al., 2011, 2009). Here, we also show the individual ROI masks (spherical mask, radius 10 mm) that were used for the LI calculations. All images were visualized with nilearn (version 0.7.0) (*Abraham et al., 2014*).

### 3.2. Distribution of hemispheric lateralization in the core system across subjects

A lateralization index was calculated for each subject and each region of the core system. Due to weak brain activity (i.e., not sufficient activated voxels in the ROI masks even at liberal statistical thresholds), an LI could not be calculated for three subjects for the STS and for two subjects for the OFA. All subsequent results are thus based on 108 LI values for the FFA, 106 LIs for the OFA and 105 LIs for the STS.

The distribution of hemispheric lateralization across the population is presented separately for right- and left-handers in Fig. 3. Hemispheric lateralization was continuously distributed across subjects for both groups, ranging from right- to left-hemispheric dominance. Particularly striking here is the high interindividual variability. For right-handers, the mean LI was −0.124 (SD = 0.490, median = −0.200) for the OFA, −0.225 (SD = 0.435, median = −0.300) for the FFA and −0.082 (SD = 0.462, median = −0.145) for the STS. For left-handers, the LI was −0.173 (SD = 0.524, median = −0.340) for the OFA, −0.121 (SD = 0.438, median = −0.220) for the FFA and −0.220 (SD = 0.508, median = −0.340) for the STS.

**Fig. 3.**
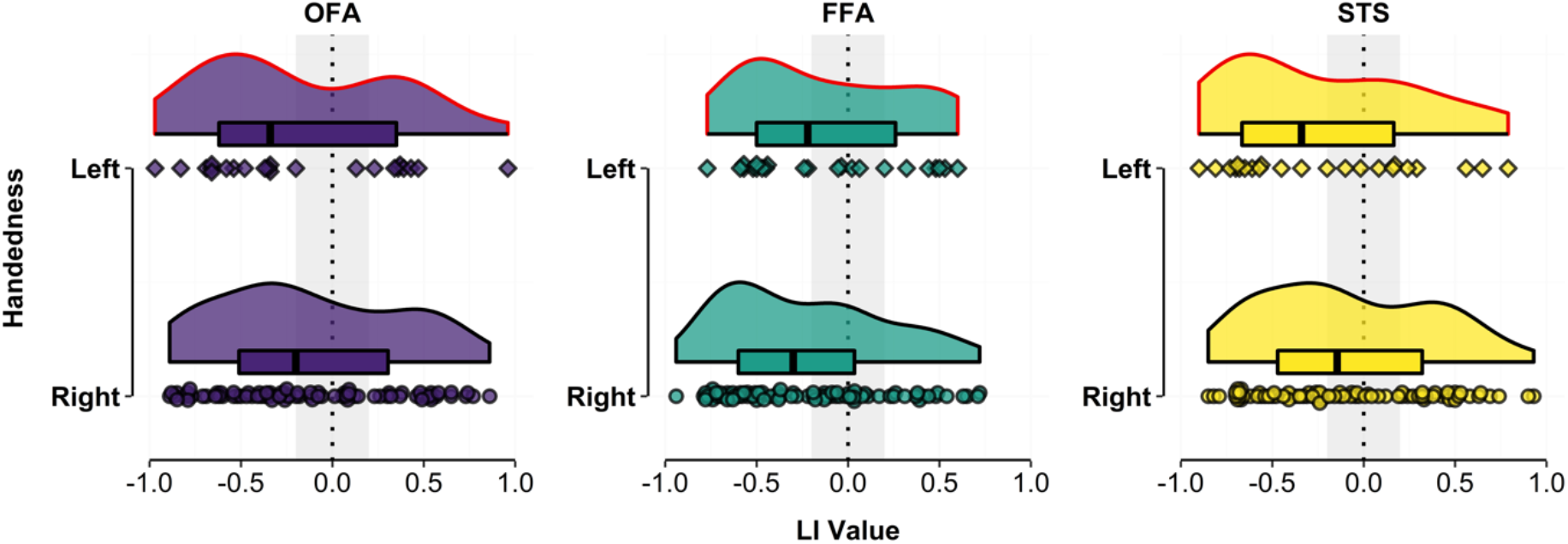
Distribution of hemispheric lateralization (described by lateralization index [LI]) for face perception across the population, separately presented for each region of the core system (OFA, FFA, STS) and handedness groups (left-handers, right-handers). Of note: negative LI values on the left side of the x-axis represent right-hemispheric dominance and positive values on the right side represent left-hemispheric dominance. Box plots with median (black vertical line) and individual data points are also plotted at the bottom of each density distribution.

All density distributions were slightly skewed to the right, indicating that overall, more subjects were right-dominant than left-dominant. Nevertheless, it is evident that there is a substantial number of subjects with bilateral or even left-hemispheric dominance. In Fig. 4 (left), we present the percentage of subjects classified as left-dominant (LI > 0.2), bilateral (|LI| ≤ 0.2) or right-dominant (LI < −0.2). In Fig. 4 (right), we additionally use a bipartite division (i.e., omit the category bilateral) and present the percentage of subjects classified as left-dominant (LI > 0.0), and right-dominant (LI < 0.0). This classification is performed both for all subjects and separately for right- and left-handers. Only about half of the subjects can be classified as right-dominant using a tripartite division, and only two-thirds of the subjects using a bipartite division. The most strongly lateralized brain region is the FFA in right-handers using bipartite division. However, also in this case 32% of subjects are not right-dominant.

**Fig. 4.**
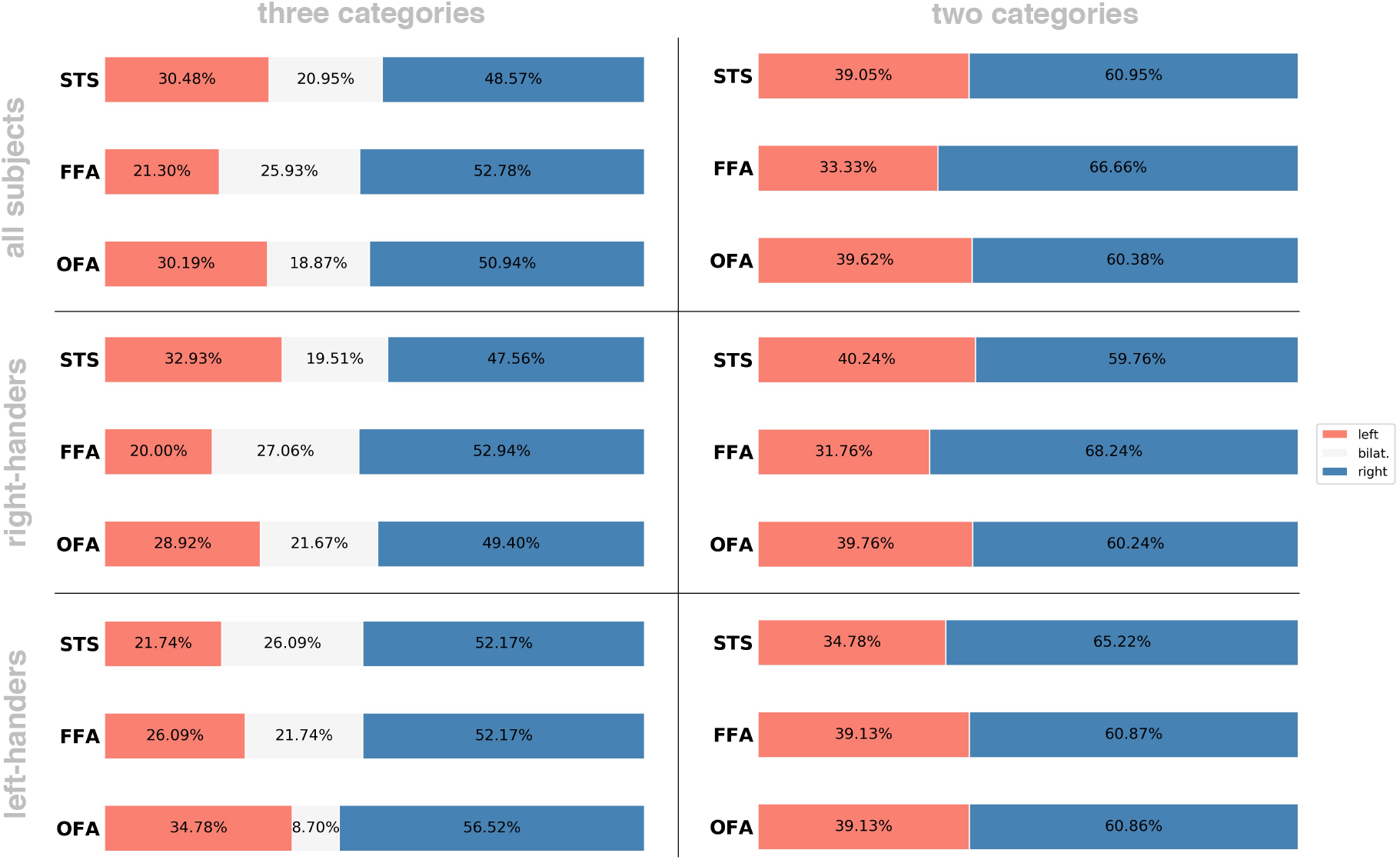
**Left column:** The percentage breakdown of hemispheric dominance in the three categories left-dominant (red), bilateral (grey) and right-dominant (blue). For all three brain areas only ~50% of the sample is right-dominant. **Right column:** Without a bilateral category, more than 60% of subjects show a right hemispheric dominance for all core system regions.

### 3.3. Effects of region, handedness and sex on hemispheric lateralization

To analyze the effects of the brain region (OFA, FFA, STS), handedness (right-handed, left-handed) and sex (male, female) on the LI, we fitted a generalized linear mixed-effects regression model. Main effects and interactions were assessed via type III Wald Chi^2^-tests. We used an alpha level of 0.05 for all statistical tests. When necessary, p-values were adjusted according to the false discovery rate.

#### Effect of brain region

Our first aim was to test the hypothesis that FFA and STS are stronger lateralized than the OFA. Our analysis, however, did not show a significant main effect of brain region (χ^2^ (2, N = 108) = 0.0875, p = 0.9572). This was also the case when we assessed models separately for right- and left-handers (see OSF, https://osf.io/s8gwd/). For right-handers, the mean LI for the OFA was −0.124 +/- 0.490, in comparison to −0.225 +/- 0.435 for the FFA and −0.082 +/- 0.462 for the STS. For left-handers, the mean LI for the OFA was −0.173 +/- 0.524, in comparison to −0.121 +/- 0.438 for the FFA and −0.220 +/- 0.508 for the STS. Thus, our data do not support the hypothesis that the OFA is on average less lateralized than the FFA and STS.

#### Effect of handedness

Our second aim was to test the hypothesis that left-handers show a reduced right-hemispheric lateralization for the FFA. Again, our data analysis did neither yield a main effect of handedness (χ^2^ (1, N = 108) = 0.0638, p = 0.8006) nor an interaction effect of handedness and brain region (χ^2^ (2, N = 108) = 4.6216, p = 0.0992). Only descriptively, the comparison of LI values for left-handers and right-handers showed a small leftward shift for the FFA (LH: −0.121 (SD = 0.438); RH: −0.225 (SD = 0.462)), while OFA (LH: −0.173 (SD = 0.524); RH: −0.124 (SD = 0.490)) and STS (LH: −0.220 (SD = 0.508); RH: −0.082 (SD = 0.462)) were even stronger right-lateralized in left-handed subjects.

We exploratively tested if handedness would only influence lateralization in combination with subjects’ sex. LI values separately for each region, handedness and sex are depicted in Fig. 5. Results indicated that left-handed men show systematic differences in lateralization compared to right-handed males and right-and left-handed females. Their FFA is bilateral with a tendency to left hemispheric dominance (LI = 0.156, SD = 0.411), while the FFA in all other subgroups is right dominant (see Table 3). An additional ordinary least squares regression model only including LI values for the FFA confirmed a significant handedness and sex interaction for the FFA (F(1, 103) = 4.7813, p = 0.031). This interaction is driven by left-handed men being significantly less right-lateralized compared to right-handed men (t(103) = 2.459, p = 0.0156).

**Fig. 5.**
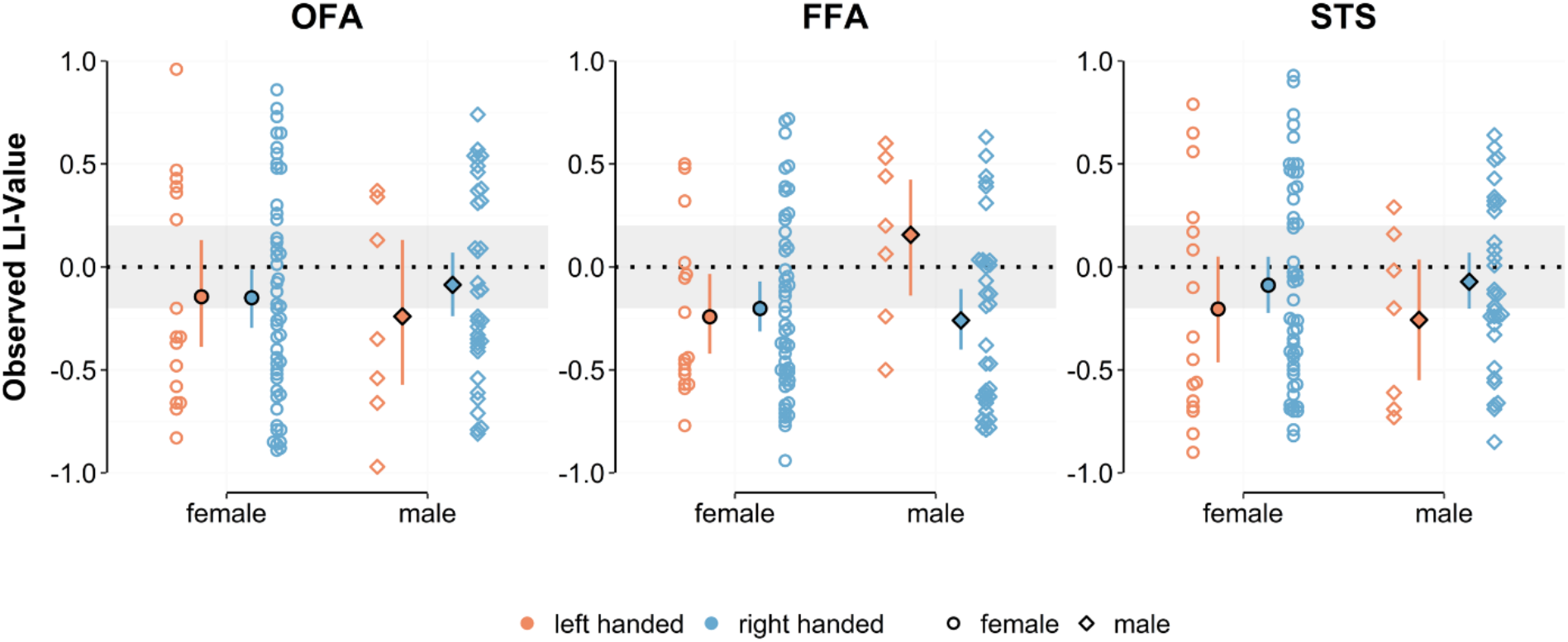
Interaction plot for brain region*handedness*sex. Mean LI values with bootstrapped confidence intervals are plotted next to the individual datapoints. Left-handed men (red diamonds) stand out as a group with a shift towards a left dominant FFA. The horizontal grey bar highlights the range for bilaterality (|LI| < 0.2).

**Table 3.** Mean (SD) LI values separate for brain region, sex and handedness.

| ROI | LH female | LH male | RH female | RH male |
| --- | --- | --- | --- | --- |
| OFA | -0.144 (0.538) | -0.240 (0.525) | -0.149 (0.512) | -0.087 (0.462) |
| FFA | -0.242 (0.402) | 0.156 (0.411) | -0.202 (0.433) | -0.259 (0.441) |
| STS | -0.204 (0.553) | -0.257 (0.422) | -0.089 (0.495) | -0.072 (0.413) |

ROI = region of interest, LH = left-handed, RH = right-handed

#### Other effects

For completeness, our analysis also did not show a main effect of sex (χ^2^ (1, N = 108) = 0.2867, p = 0.5924), nor a three-way interaction of brain region, handedness and sex (χ^2^ (2, N = 108) = 3.8174, p = 0.1483) or a two-way interactions of brain region and sex (χ^2^ (2, N = 108) = 1.1249, p = 0.5698) or of sex and handedness (χ^2^ (1, N = 108) = 0.1821, p = 0.6695).

## 4. DISCUSSION

Hemispheric lateralization is a fundamental principle of brain organization in humans and many other species (Esteves et al., 2020; Güntürkün et al., 2020; Karolis et al., 2019). Asymmetry rather than symmetry seems to be ubiquitously present in brain anatomy and function (Esteves et al., 2020). Theoretical advantages of hemispheric asymmetries include parallel processing of complementary information, maximization of available space, higher proficiency and processing speed and decreased inter-hemispheric competition (Esteves et al., 2020; Güntürkün et al., 2020). Lateralization patterns are highly flexible and can be modulated by various factors like handedness, sex, age, genetic factors, hormonal influences, or disease (Toga and Thompson, 2003).

In face perception research it is generally accepted that the right hemisphere is playing the dominant role (Duchaine and Yovel, 2015; Rossion and Lochy, 2021). This observation is based on ample evidence accumulated over the last decades with lesion patients, brain stimulation studies, and behavioral experiments. Prosopagnosia is, for instance, mostly caused by right-hemispheric lesion, only seldom by unilateral damage to the left hemisphere (for an overview, cf. Bukowski et al., 2013). In contrast, fMRI studies comparing left- and right-hemispheric activation in homologous face sensitive areas have painted a picture of large variability between studies and amongst individual subjects ranging from clear right dominance over bilaterality to individuals with left hemispheric dominance. Notably, most of these studies are based on rather small sample sizes often deliberately excluding left-handers (Willems et al., 2014) or only assessing the FFA. Thus, the present fMRI study aimed to systematically determine hemispheric lateralization of all core system regions in a relatively large cohort (N = 108) of healthy right- and left-handers. We were particularly interested in the variability of hemispheric lateralization across subjects and explored how many subjects can be classified as right-dominant based on the fMRI activation pattern. We further intended to determine lateralization differences between different regions of the core system and to assess the influence of handedness and sex on the lateralization pattern.

### 4.1. Interindividual variabiliy of hemispheric lateralization

Hemispheric lateralization was continuously distributed across subjects, ranging from strong left-to strong right-hemispheric dominance both in right- and left-handers. Depending on the specific region, the mean LI ranged from −0.082 to −0.225. At the group level, the degree of hemispheric lateralization of the core system of face perception network could thus be characterized as “bilateral to weakly right-dominant”. Right-hemispheric lateralization was strongest for the FFA in right-handers (mean LI = −0.225), but the degree of lateralization had to be characterized as “weak” in comparison to other cognitive brain functions such as language or spatial attention. The hemispheric lateralization of the language network is for instance characterized by LIs that are typically larger than 0.5 (e.g., Somers et al., 2011).

Particularly striking was the high interindividual variability. While the LI could by definition only vary between −1 and 1, the standard deviation ranged from 0.435 to 0.529, expressing a high level of dispersion around the mean. Only approximately 50% of subjects could be classified as right-hemispheric dominant when applying the widely used criterion for bilaterality (|LI| < 0.2). Even if we classified all subjects with LI values below 0.0 as right-hemispheric dominant (i.e., omit the category “bilateral”), still about one third of subjects had a dominant left hemisphere.

The high interindividual variability and the large number of subjects who do not show a clear right-hemispheric dominance is also evident from other studies (e.g., Canário et al., 2020; Davies-Thompson et al., 2016; De Winter et al., 2015; Johnstone et al., 2020), with up to 45% of subjects being not right-hemispheric dominant for face perception. This issue is usually not explicitly discussed. It is, however, abundantly obvious from the presented data. It would thus be too simple to argue that the observed inter-individual variability of LI values is a specific feature of our present study or the specific task.

Taken together, data both from the present and previous studies provide only limited support for a clear right-hemispheric dominance of the face perception network based on fMRI activation patterns alone. If one assessed the fMRI activation pattern for face perception independent from previous expectations derived from the results of other modalities (e.g., lesion studies), one would most likely not be inclined to characterize the network as right-hemispheric dominant. Instead, one would rather term it “bilateral with a slight tendency towards right-dominance at the group level and a large interindividual variability”.

To avoid misunderstandings, we would like to explicitly state that we do not question the right-hemispheric dominance of the face perception network per se. Lesion studies, for instance, clearly show, as mentioned several times previously, that damage to the right hemisphere is typically associated with more obvious behavioral deficits than damage to the left hemisphere (Duchaine and Yovel, 2015; Rossion and Lochy, 2021). Using this criterion, the face perception network can be characterized as right-dominant. We only question that the fMRI activation pattern associated with face perception should be characterized as “typically right-hemispheric dominant”.

At this point, it is important to understand how the discrepancy between the high percentage of subjects classified as right-hemispheric dominant based on lesion data and the much lower ratio of individuals classified as right-hemispheric dominant based on fMRI data arises. A possible explanation is that the hemispheric lateralization derived from lesion data and the lateralization derived from fMRI data simply reflect different aspects of face perception. In this case, they do not necessarily have to be strongly correlated.

From lesion data, one can infer that specific brain regions are necessary for the execution of certain cognitive functions (or at least specific aspects of these functions). The right-hemispheric dominance of face perception, as determined by lesion studies, thus describes that specific aspects of face perception (e.g., identity processing) are typically (i.e., in most subjects) lateralized to the right hemisphere. From fMRI data, one can infer that face perception is associated with a bilateral network. Despite decades of research, however, we still lack a precise characterization of the functional differences between the left- and right-hemispheric homologues of the face perception network. Various hypotheses have been proposed to characterize their different functional profiles. They typically describe the processing style of the two hemispheres in a dichotomous fashion (e.g., left high vs. right low spatial frequencies (Sergent, 1982), left analytic vs. right holistic (Bradshaw and Nettleton, 1981; Rossion et al., 2000); left proactive vs. right reactive; left local vs. right global; see also Rossion and Lochy (2021) and Dien (2008, 2009) for an overview). Interindividual differences in the fMRI lateralization pattern might thus be associated with individual differences in the processing styles and might be less related to the probability of showing behavioral deficits after lesions to either hemisphere.

The challenge of future research will be to delineate potential factors driving hemispheric lateralization of fMRI activation patterns not only at the group level, but also in individual subjects. In other words, while decades of research investigated the underlying mechanisms of a right dominant face perception network (e.g., neural competition hypothesis with language areas, see Behrmann and Plaut, 2020, 2015; Dehaene et al., 2015, 2010; Rossion and Lochy, 2021) one should now address the question why – based on fMRI – roughly 50% of subjects do not show this so called “typical” pattern. Consequently, if it is totally normal for half of the population to present an atypical lateralization pattern, this brings up the questions whether lateralization patterns of individual subjects’ matter at all in terms of healthy brain function or disease. For example, individuals with left dominant face sensitive areas could potentially rely on other processing strategies than individuals with a right hemispheric dominant face network. Furthermore, a specific lateralization pattern could also be favorable for specific face perception tasks or even indicative of certain psychiatric diseases.

### 4.2. Effects of region, handedness, and sex on hemispheric lateralization

The second aim of our study was to investigate whether there are lateralization differences between different regions of the core system and to assess the influence of handedness and sex on the lateralization pattern. Our data did neither show a significant main effect of region, handedness, or sex nor any interaction. However, in an exploratory analysis of the FFA lateralization alone, we showed that handedness in combination with sex has an influence on hemispheric dominance, with left-handed men having a significantly more left-lateralized activation pattern compared to all other subgroups. These results are discussed in the following.

#### Are core system regions differentially lateralized?

Faces are generally processed in direction from posterior (e.g., OFA) to anterior (e.g., ATL, IFG) brain regions. This is also reflected by simultaneous recordings of EEG and fMRI responses to faces, showing that OFA activation is correlated with early event-related potentials about 110 ms after stimulus onset, while activations in the temporal lobe (i.e., FFA and STS) are highly correlated with the later face-sensitive N170 component (Sadeh et al., 2010). Whether these latency effects also translate to increased lateralization in posterior to anterior direction within the core system remains largely unknown. Initial evidence in this regard was provided by Rossion and colleagues (2012) for right-handed subjects. They calculated the percentage of activated voxels in the right hemisphere and found no significant lateralization for the OFA (61%), but significant right-hemispheric dominance for the FFA (71%) and STS (77%).

Against our expectations we did not find a significant main effect of brain region (χ^2^ (2, N = 108) = 0.0875, p = 0.9572). Our data thus did not support an increase in hemispheric dominance from posterior (i.e., OFA) to anterior (i.e., FFA and STS) regions. The activation pattern of the OFA was found to be bilateral in both right- (LI = −0.124 ± 0.490) and left-handed subjects (LI = −0.173 ± 0.524). Even the brain region showing the most lateralized activity, the FFA in right-handers (LI = −0.225 ± 0.435), was only slightly stronger lateralized.

Remarkably, a comparison of lateralization results found in the fMRI literature revealed one obvious pattern: inconsistency for all three face sensitive areas. For each area there are several studies showing clear right-hemispheric lateralization (e.g., Frässle et al., 2016c; Ishai et al., 2005; Rhodes et al., 2004; Rossion et al., 2012), while others found inconclusive results (e.g., Haxby et al., 1999; Yovel et al., 2008) or even bilateral activity without a significant lateralization effect (Canário et al., 2020; De Winter et al., 2015; Ishai et al., 2002). In summary and in line with our results, face sensitive areas in the core system only show a gentle tendency towards the right hemisphere. Furthermore this tendency is just a group effect with limited implications for individuals.

#### Does handedness influence lateralization?

Approximately 90% of humans are right-handers. This proportion has remained relatively stable over the past 5000 years (Coren and Porac, 1977). From language research it is well known that handedness can influence hemispheric dominance. While 96% of right-handed subjects show a left hemispheric langugage dominance, this value is reduced to 73% in left-handers (Knecht et al., 2000b). Thus, to rule out handedness effects, face perception research has predominantly focused on lateralization patterns of right-handers (Rossion et al., 2012), deliberately neglecting the investigation of left-handers (Willems et al., 2014).

In the last decade, however, a handful of research groups have specifically recruited left-handers and detected a significant reduction in FFA lateralization compared to right-handers (Bukowski et al., 2013; Frässle et al., 2016a; Willems et al., 2010; see also Badzakova-Trajkov et al., 2010 for lateralization in the whole temporal lobe). Precisely, neither the percentage of activated voxels (Bukowski et al., 2013; Willems et al., 2010) nor the activation strength (Frässle et al., 2016a) was found to be significantly lateralized in left-handers. On the other hand, OFA (see Bukowski et al., 2013; Frässle et al., 2016a) and STS (Bukowski et al., 2013) were right dominant and not significantly different from right-handers. Bukowski et al. (2013) speculated that the observed bilaterality of the FFA is not reflecting a true bilaterality but could instead be caused by broader variations of LI results amongst left-handers.

However, in our sample we neither found a significantly reduced right-hemispheric dominance in left-handers (main effect handedness: χ^2^ (1, N = 108) = 0.0638, p = 0.8007) nor an increased variability of LI values (e.g., left-handed: SD FFA = 0.438, right-handed: SD FFA = 0.435). Only the interaction of handedness and brain region indicated a tendency towards the expected leftward shift of FFA lateralization in left-handed subjects (χ^2^ (2, N = 108) = 4.6216, p = 0.0992). Notably, all of these studies including ours assessed rather small cohorts of left-handed subjects (n < 33) and used different paradigms and analyses methods. Therefore, large scale studies with left-handed subjects are needed to discern whether the handedness effects found in previous studies are genuine to the left-handed population or rather caused by a sampling bias or specific methodological decisions.

One possible cause of variability amongst left-handers might be their language dominance. For example, two studies conducted by Gerrits and colleagues (2019, 2020) both revealed a reduced right-hemispheric dominance in left-handers with atypical right language dominance. Left-handers with typical left language dominance on the other hand were not distinguishable from right-handers. Lastly, one could also think of sex as a potential source of variance amongst the left-handed population (as discussed below).

#### Are men and women differentially lateralized?

Sex differences in hemispheric asymmetry and their implications on cognitive functioning like mental rotation abilities or verbal skills have been a matter of debate for many decades (Hirnstein et al., 2019). For example, male brains have repeatedly shown to be more lateralized than female brains, albeit with a very small effect size (Hirnstein et al., 2019). Furthermore, myriads of studies reported a male advantage in mental rotation tasks (Zell et al., 2015), while women often outperform men in verbal tasks (Lowe et al., 2003).

However, a causal link between cognitive performance and hemispheric asymmetry related to sex is still missing. To our knowledge no differences between males and females in hemispheric lateralization in the face perception network have been reported so far (e.g., see Rossion et al., 2012 who found no evidence for female vs. male difference in their lateralization pattern of OFA, FFA or STS). In line with these observations the present study did not find a significant main effect of sex.

Interestingly, when exploratively combining sex and handedness effects specifically in the FFA, one group of subjects stood out in particular: left-handed men (see Fig. 5). Compared to right-handed men, left-handed man showed a significant left-ward shift of FFA lateralization (mean LI left-handers = 0.149, SD = 0.450). This observation is based on the seven left-handed men included in our sample, rendering the need for larger samples to substantiate this finding.

### 4.3. Limitations

Last, we would like to point out potential limitations of our study. The LI is influenced by a myriad of factors. Being based on fMRI activation patterns, it obviously depends on the chosen paradigm, fMRI acquisition parameters, preprocessing pipelines, the choice of an activity measure and the definition of suitable regions of interest (for an overview, cf. Jansen et al., 2006). Since the regions of the core system are in close anatomical proximity, different from for example Broca’s and Wernicke’s area in the language network, we have spent in particular some effort to localize these regions in a meaningful way in individual subjects (Fig. 1). Further studies, however, should investigate in more detail the influence of specific analysis parameters on hemispheric lateralization. This will also help to make results from different studies more comparable.

### 4.4. Conclusions

In summary, our fMRI data did not support a clear right-hemispheric dominance of the face perception network. On average, brain activity was stronger in right-hemispheric brain regions than in their left-hemispheric homologues. This asymmetry was, however, only weakly pronounced in comparison to other lateralized brain functions (such as language and spatial attention) and strongly varied between individuals. Only half of the subjects in the present study could be classified as right-hemispheric dominant. Instead of calling the fMRI activation pattern right-dominant, as is done in most studies, it might be more suitable to characterize it as “bilateral, with a slight tendency towards right-dominance at the group level and a large interindividual variability”. Our findings ultimately question the dogma that the face perception network – as measured with fMRI – is typically right lateralized. To put it bluntly, how can something be typical if it only applies to half of the population? This would be equally precise as terming the world population typically female.

## Credit authorship contribution statement

**Ina Thome:** Conceptualization, Data curation, Formal analysis, Investigation, Methodology, Project administration, Software, Visualization, Writing – original draft, Writing – review & editing. **José C. García Alanis:** Formal analysis, Methodology, Visualization, Writing – review & editing. **Jannika Volk:** Data curation, Formal analysis. **Christoph Vogelbacher:** Data curation, Writing – review & editing. **Olaf Steinsträter:** Software. **Andreas Jansen:** Conceptualization, Supervision, Resources, Funding acquisition, Writing – review & editing.

## Aknowledgements

This work was supported by the German Research Council (Deutsche Forschungsgemeinschaft, DFG, Grant no. JA 1890/11-1) and the University Medical Center Giessen and Marburg (UKGM) (Grant no. 7/2019-MR).

We would like to thank Stefan Frässle, Anna Rysop, Peer Herholz, Verena Schuster, Mechthild Wallnig, Rita Werner, and Jens Sommer for the setup of the experiment and raw data acquisition.

## Data/code availability statement and supplementary material

Supplementary material as well as data and code used in this study are publicly available on the Open Science Framework at https://osf.io/s8gwd/.

